# Protein Design with StructureGPT: a Deep Learning Model for Protein Structure-to-Sequence Translation

**DOI:** 10.1101/2024.06.03.597105

**Authors:** Nicanor Zalba, Pablo Ursua-Medrano, Humberto Bustince

**Affiliations:** Computational Protein Design, InsAIght Research Center, Monasterio de Tulebras 2, Office 5, 31011, Navarra, Spain; Statistitcs, Informatics and Mathematics, Public University of Navarra, Cataluña Avenue, 31006, Navarra, Spain

**Keywords:** deep learning, structure-to-sequence translation, protein design, natural language, generative transformer, protein solubility, protein stability

## Abstract

**Motivation:** Protein design, crucial for understanding and engineering protein functionalities, has traditionally been challenged by the reverse translation of complex protein tertiary structures into sequences. Existing computational tools have focused predominantly on sequence-to-structure predictions, with less attention given to structure-to-sequence processes. Our research introduces StructureGPT, a novel deep learning model that employs advanced natural language processing techniques to translate complex protein tertiary structures into their corresponding amino acid sequences. This model addresses critical gaps in protein engineering, particularly improving solubility and stability, which are essential for pharmaceutical development and industrial applications.

**Results:** StructureGPT demonstrates the capability to autoregressively generate amino acid sequences from detailed structural inputs, enhancing the design of proteins with specific functionalities. By leveraging the linguistic parallels between protein structures and human language, our model not only predicts sequences with high accuracy but also suggests modifications that could lead to improved protein properties. The application of StructureGPT in multiple protein design tasks showcases its utility in various biomedical and biotechnological contexts.

**Availability:** The source code for StructureGPT is freely available at https://github.com/StructureGPT DOI: 10.5281/zenodo.11065607

## 1 Introduction

Proteins are essential to myriad biological processes and offer vast potential for applications across research, industry, and pharmaceuticals due to their functional versatility. The advent of computational models has significantly advanced our ability to design and manipulate these molecules for specific applications. One notable example, AlphaFold2, introduced by Jumper et al. in 2020, has revolutionized the field by predicting protein structures from amino acid sequences with unprecedented accuracy (1). However, the challenge of translating complex protein tertiary structures back into sequences remains less explored and equally critical, especially for understanding and engineering protein functionality.

Our contribution to this evolving field is StructureGPT, a compact deep learning model trained to perform this reverse translation. By employing techniques from NLP, StructureGPT interprets the ‘language’ of protein structures, enabling it to generate corresponding amino acid sequences. This capability allows for the application of StructureGPT across multiple protein design tasks, thus enhancing its utility in various biomedical and biotechnological contexts (5; 6; 2; 3; 4).

In this work, we demonstrate how StructureGPT not only supports the generation of multiple amino acid sequences from a given protein’s structure but also optimizes sequences for specific functional properties such as solubility and stability. This is accomplished through autoregressive generation conditioned on detailed structural information, showcasing the model’s ability to incorporate absent structural elements during sequence prediction (7; 8; 9; 10). Our approach marks a significant step forward in the field of protein engineering, offering a new tool for researchers to create novel protein variants with tailored functionalities, ultimately leading to innovations in healthcare and environmental sustainability (11; 12; 13; 15; 14).

## 2 Material and Methods

### 2.1 Structure dataset

For model training, two independent datasets were constructed using different subsets of tertiary structures predicted by AlphaFold2 for amino acid sequences annotated in the Swiss-Prot (40) and UniProt (39) databases. In both cases, the structures are available for download at https://alphafold.ebi.ac.uk/download.

The first dataset, named SwissProtDB, was formed with tertiary structures of less than 1200 amino acids corresponding to annotated sequences in the Swiss-Prot database. 10% of these structures, randomly selected, were used to compose the validation and test sets. Consequently, the training and validation sets consisted of 469 635 and 26 256 structures, respectively.

The second dataset, named AlphaFoldDB 1M, was constructed with tertiary structures of less than 1700 amino acids corresponding to the proteomes of the organisms Homo sapiens, Mus musculus, and Escherichia coli. In total, around 1 million structures were grouped, of which approximately 10%, randomly selected, were allocated to form the validation and test sets. Thus, the final training and validation datasets contained 1 019 584 and 54 298 structures, respectively.

### 2.2 Data encoding

The Cartesian coordinates of the atoms forming each amino acid were grouped to form a vector. Each type of atom was assigned one of the 38 classes represented in Table S1. Similarly, three fixed positions within the vector were assigned to each atomic class, one for each Cartesian coordinate. This process resulted in a final vector of 114 components for each amino acid. These vectors plays a role analogous to that of a word vectors in an NLP problem. Finally, the representation of the tertiary structure of each complete protein was obtained by concatenating the vectors representing individual amino acids. As a first step, centers of mass for each amino acid in the structure of utrophin were calculated. Then, distances between centers of mass, as well as between their corresponding PCA points were determined. After that, linear and monotonic correlation between distances was assessed through Pearson and Spearman correlation analysis, respectively. The centers of mass were used as descriptors of the structural context since all the forces acting on the atoms of an amino acid can be assumed to be applied at that point. This also assumes rigid behavior of the amino acids against the forces acting on them. Centers of mass, Euclidean distances between PCA points and between centers of mass were computed using the BioPython library (31).

To evaluate the linear and monotonic relationships between the distances from PCA points to the distances from centers of mass, Pearson and Spearman correlation analyses were performed, respectively. These statistical analyses were facilitated by the SciPy library (34).

### 2.3 Model architecture

The representation of the structure we use in this work significantly differs from the one commonly given to words in classical language problems. In such problems, each word or word fragment is usually identified with an index within a vocabulary, and each index corresponds to a word vector during the embedding process. Typically, the number of components in embedding vectors arises from a compromise between computational efficiency and the vector’s ability to capture semantic information about the word. In our case, the initial dimensionality of the vectors representing each amino acid does not follow this criterion but is determined by the need to capture structural information as faithfully as possible. However, to ensure that these vectors capture the maximum amount of information, a projection to a higher dimensionality was carried out. To achieve this, it was necessary to introduce a modification in the architecture of the classical Transformer encoder-decoder (22), where the word embedding block was replaced by a linear layer (see Figure S1).

One way to solve a translation problem between two languages is to liken it to a multiclass classification problem. Thus, one of the tasks that a language model must perform is to identify each word with a class. Natural languages generally consist of many words, so many classes are usually required to faithfully capture them. However, for the structure-sequence translation problem, only 20 different classes are needed to represent the natural amino acids found in proteins. According to these ideas, StructureGPT was trained to solve a multiclass classification problem in which the sequence language is captured in 20 different classes. However, since the structure of a protein can include multiple chains, chain and sequence start and end symbols were also included in the sequence language.

### 2.4 Model training

The implementation of the Transformer encoder-decoder for this work consisted of 6 encoder and 6 decoder blocks, each with 8 attention heads, an embedding dimension of 512, and a feedforward dimension of 2048.

This architecture was independently trained with the two datasets described in a previous sections. Although both training processes were conducted under similar conditions, the difference in the data used each time necessitated introducing some variations between them. In both cases, the AdamW optimizer was used (*β*_1_=0.9, *β*_2_=0.98, *ϵ*=10^−9^), parameters were initialized using the Xavier uniform algorithm, and CosineAnnealingLR was applied as a scheduler to reduce the learning rate from 10^−4^ to 6 *·* 10^−5^.

Both trainings were conducted on 2 NVIDIA A100 with 40 GB of VRAM, using a batch size of 4 examples. The first training was completed in 37 epochs (approximately 9 days), while the second training required 70 epochs (approximately 2 months).

### 2.5 Stability modulation of human frataxin

To generate the different mutational variants of the mature form of human frataxin (amino acids 90 to 210), 14 mutational hotspots were previously identified through a conservation analysis similar to the one described in (23). The initial sequence alignment, consisting of 2776 sequences, was performed against the UniRef90 database (database date 2024-02) with the help of the *jackhammer* (33) algorithm (flags –F1 0.0005 –F2 5e-05 –F3 5e-07 –incE 0.0001 -E 0.0001 –N 1). Then, for simplicity, only a multiple sequence alignment (MSA) with sequences that had at least 80% identity with human frataxin (MSA80) was constructed, resulting in an alignment formed by 227 sequences. Through a conservation analysis on the MSA80, positions 101, 116, 117, 120, 121, 179, 187, 188, 192, 195, 197, 202, 205, and 208 of the WT frataxin sequence were identified as mutational hotspots. Next, to generate the different variants of the frataxin sequence, StructureGPT was asked to translate into sequence the structure of human frataxin collected in the Protein Data Bank (PDB ID 1EKG), considering for the amino acids located at hotspots two different atomic classes. Each of the sequences obtained corresponds to one of the possible combinations that result from considering two different amino acid classes for each of the hotspots. As a result of this procedure, 16,384 sequences were finally obtained.

Subsequently, with the help of software developed for this purpose, those sequences with the same length as the sequence of the mature form of frataxin and that included mutations only in one or several of the mutational hotspots were filtered. In this way, 56 different sequences were identified (Figure S4).

For each of the selected mutant sequences, the three-dimensional structure was calculated using AlphaFold Colab, available at https://colab.research.alphafold (num relax: 0, template mode: none, msa mode: mmseq2-uniref-env, pair mode: unpaired paired, model type: auto, num recycles: 3, recycle early stop tolerance: auto, relax max iterations: 200, pairing strategy: greedy, max msa: auto, num seeds: 1). For each mutant, the structure for which AlphaFold offered the highest predicted local distance difference test (pLDDT) was chosen; these structures were then used to calculate the Δ*G*_*F*_ using FoldX (27; 25). Finally, the ΔΔ*G*_*F*_ were calculated as the difference between the Δ*G*_*F*_ of each mutant and the corresponding Δ*G*_*F*_ value for frataxin in its mature form.

### 2.6 Solubility modulation of rhGM-CSF

Since the annotated structures for rhGM-CSF in the PDB (IDs 1CSG and 2GMF) correspond to dimers and StructureGPT can only process structures of monomeric proteins, the first step was to calculate the structure of the protein in its monomeric form using AlphaFold Colab from the sequence of chain A in 1CSG (num relax: 0, template mode: none, msa mode: mmseq2 uniref env, pair mode: unpaired paired, model type: auto, num recycles: 3, recycle early stop tolerance: auto, relax max iterations: 200, pairing strategy: greedy, max msa: auto, num seeds: 1, the structure with the highest overall pLDDT was chosen). Simultaneously, a conservation analysis similar to that carried out for human frataxin was performed (jackhammer flags –F1 0.0005 –F2 5e-05 –F3 5e-07 –incE 0.0001 -E 0.0001 –N 1, MSA depth: 122 sequences, MSA80 depth: 6 sequences). Although glutamine 95 and isoleucine 96 were conserved at 70%, and LEU111 was conserved at 80% (Figure S6), considering the low depth of the MSA80 and that neither of the three amino acids are essential for the protein’s function (28; 29), they were considered as mutational hotspots. StructureGPT was then requested to translate the structure into sequence considering 4 amino acid classes for each of the 2 positions. Out of the 256 resulting sequences, 156 had the same length as the original sequence, and of these, only 72 included mutations exclusively at the desired points. Finally, with the help of the online version of CamSol https://www-cohsoftware.ch.cam.ac.uk/, intrinsic solubility profiles corrected by structure were calculated for the different mutants and for the WT form of the protein (pH 7, patch radius: 10).

### 2.7 Inpainting analysis for human ATP synthase

For the generation of the two variants of the ATP synthase subunit A (UniProt ID A0A023I7V4), an analysis was conducted where fragments of the structure were sequentially removed, and StructureGPT was requested to translate the remaining structure fragments into sequences until total information replenishment was achieved. Through this methodology, information restoration was accomplished for 42 out of the 40 amino acids removed in the structure, positioned between valine 158 and leucine 198 of the WT sequence. Subsequently, StructureGPT was instructed to introduce mutations at these 42 reinstated positions in the resulting sequence using the mutational decode strategy (considering 3 amino acid classes for each position). For computational efficiency reasons, only a subset of the mutant variants, consisting of 15 000 sequences were calculated. Then, variants with the same length as the original sequence were filtered, thus obtaining 4 different variants. The structures of all variants were computed with the assistance of AlphaFold Colab (num relax: 0, template mode: none, msa mode: mmseq2-uniref-env, pair mode: unpaired paired, model type: auto, num recycles: 3, recycle early stop tolerance: auto, relax max iterations: 200, pairing strategy: greedy, max msa: auto, num seeds: 1). For both variants, structures with the highest pLDDT were chosen for the structure alignment analysis.

Sequence alignments of the human ATP synthase and its two inpainted variants were performed using the Jackhammer tool with specific parameters set to optimize sensitivity and specificity (flags –F1 0.0005 –F2 5e-05 –F3 5e-07 –incE 1e-06 -E 1e-06 -N 1). The alignment date for the UniRef90 database was February 2024. The multiple sequence alignment (MSA) included 20,111 sequences for the human ATP synthase, and 13 906 and 14 089 sequences for variants 1 and 4, respectively. Statistical comparisons between these sequences were conducted using a Mann-Whitney U analysis, which was implemented in custom-developed Python software to assess differences in sequence identity distributions.

RMSD, GDT-TS, GDT-HA4 and lDDT values for the pair alignment of the structures of ATP synthase, variant 1 and variant 4 were computed with the PyMol software (35), BioPython library (31) and in-house python software. During the calculations, the description of these metrics provided in (37; 38; 36) was followed.

## 3 Results

### 3.1 The atom encoding

The first step for the structure-to-sequence translation is to organize the atomic coordinates into a novel encoding, which we term ‘atom encoding’ (see Materials and Methods). This approach aims to preserve the contextual information of the tertiary structure in a way similar to how word vectors preserve the semantic information.

Figure 2 presents both a qualitative (Figures 2A, 2C, 2D, and 2E) and a quantitative (Figure 2B) analysis of the ability of atom encoding vectors to capture the structural context information contained in the Cartesian coordinates of the atoms that constitute utrophin (UniProt ID A0A0A0MSM3). Figure 2A displays the protein structure with polar contacts between amino acids highlighted as black dashed lines and the numbering of some of the amino acids. Figures 2C, 2D, and 2E show the points obtained from the atom encoding vectors through a principal component analysis (PCA) (24), along with the numbering that indicates which amino acid of the structure corresponds to which point. Notably, PCA allows to reduce the dimensionality of the atom encoding vectors from 114 to 2 while retaining most of the structural context information they contain. In Figure 2C, the points were obtained directly from the atom encoding vectors, whereas in Figure 2D, the atom encoding vectors were subjected to random masking of the Cartesian coordinates of some atomic classes, and in Figure 2E, the Cartesian coordinates of the atom encoding vectors were replaced by 1s as a preliminary step to PCA. It can be seen how points close in the plane correspond to amino acids close in the structure, while this proximity relationship becomes distorted when applying atomic class masking, or even lost when applying Cartesian coordinate masking.

**Fig. 1.**
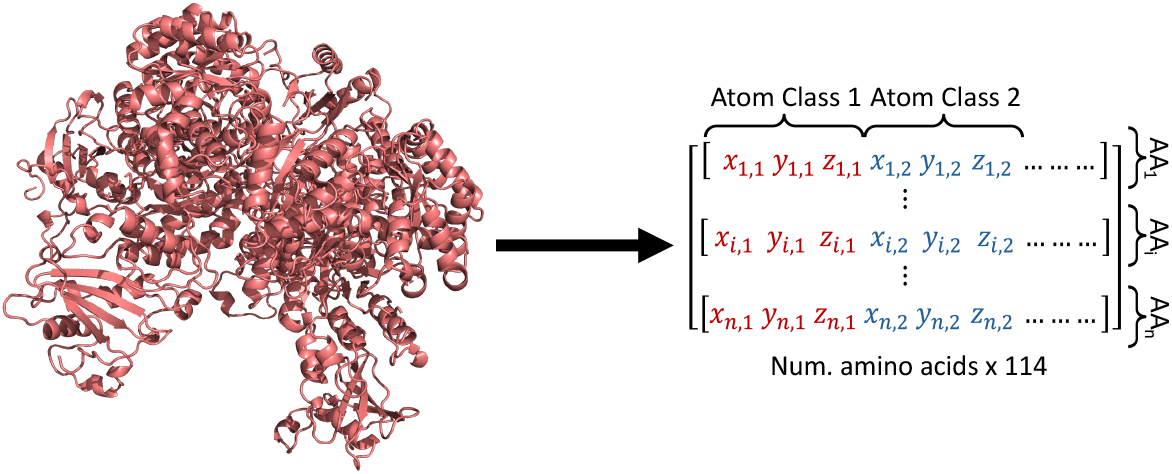
Encoding of structural information. The Cartesian coordinates are reorganized so that each amino acid corresponds to an encoding vector. Each of these vectors contains, arranged in fixed positions, the Cartesian coordinates of all the atoms that constitute it. (Right) The coordinate triples of atomic classes 1 and 2 are colored in red and blue, respectively, as an example.

**Fig. 2.**
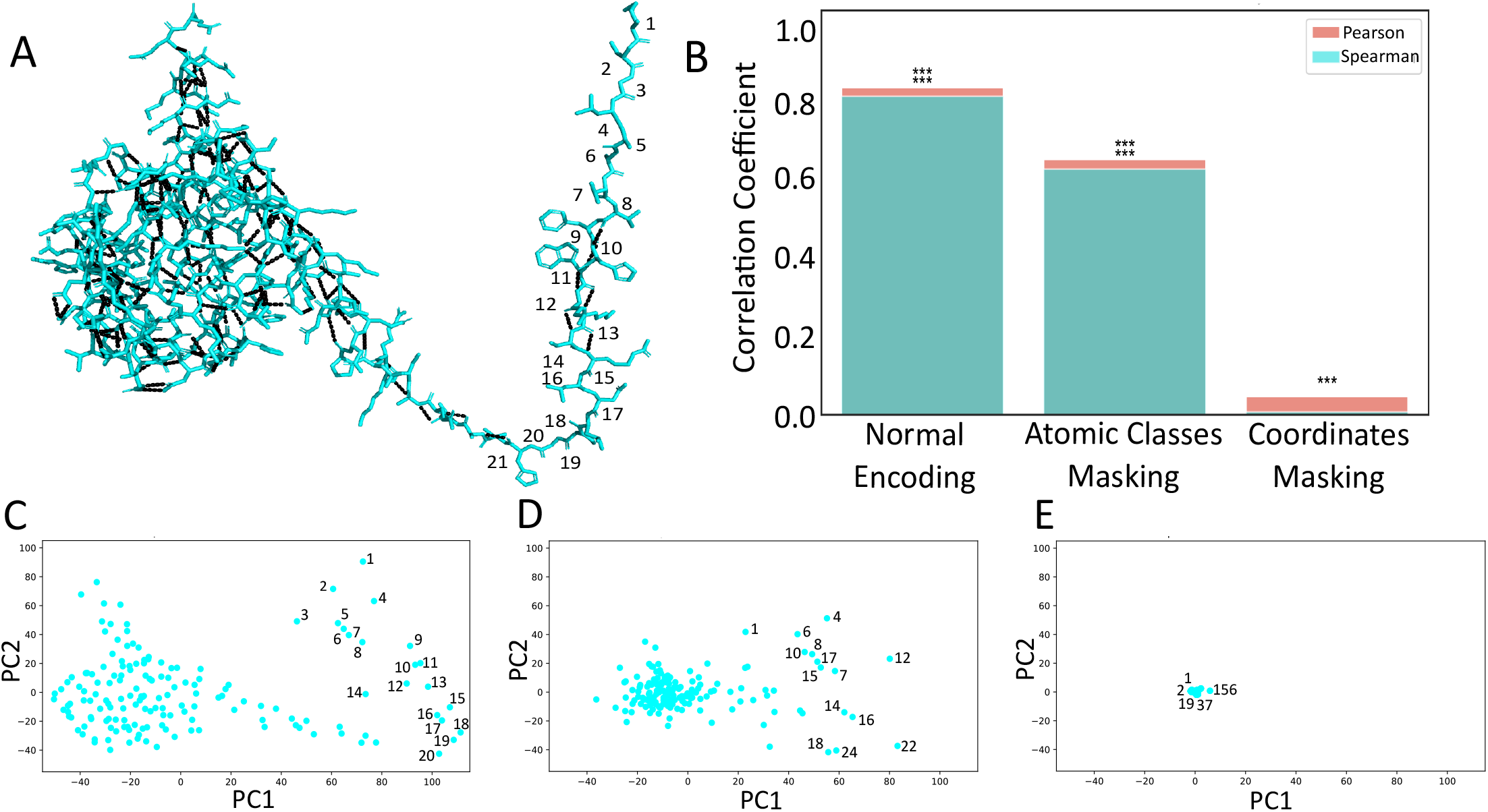
Qualitative and quantitative analysis of atom encoding properties. (A) Structure of utrophine. Polar contacts are highlighted by dashed black lines. Amino acids from 1 to 21 are numbered over the structure. (B) Statistical analysis for Pearson and Spearman correlation coefficients (from left to right) using normal encoding, with random masking of atomic classes, and with coordinate masking. Statistical significance is marked with asterisks as follows: *** for p-value *<*0.001. The tests were significant for Pearson coefficients in all three cases, and for Spearman coefficients with normal encoding and with atomic class masking. (C), (D), and (E) show the PCA points obtained from the encoding vectors. Some of the points are identified with the number of the amino acid they originate from. (C) PCA points obtained from atom encoding vectors. Points close in the plane correspond to amino acids close in the protein structure. (D) PCA points obtained from atom encoding vectors with random masking of atomic classes. The proximity relationship between plane points representing amino acids close in the structure is distorted by the masking. (E) PCA points obtained from atom encoding vectors with replacement of Cartesian coordinates by 1s in the vectors. The proximity relationship is destroyed by the coordinate masking.

To quantify how the proximity relationship of amino acids in the structure correlates with the proximity relationship of points obtained through PCA, Pearson and Spearman correlation analyses were conducted (Figure 2B). It can be observed how the Pearson correlations were statistically significant (p-value <0.001) in all three cases shown (Figure 2B), while the Spearman correlations were statistically significant (p-value <0.01) only in the first two cases. This result reinforces the idea conveyed by the quantitative analysis that the structural context captured by the atomic coordinates is preserved in the atom encoding even though the nature of both representations of the structure is very different.

### 3.2 Model training

To tackle the task of structure-sequence translation through a supervised learning approach, a training dataset consisting of protein tertiary structures labeled with their corresponding amino acid sequences was constructed (see Material and Methods). A potential source of such data could be experimentally resolved structures collected in the Protein Data Bank (30) (PDB). However, a drawback of these data is that different structures have been solved using different criteria and methodologies. This could lead to biases in the model due to potential disparities among structures. To address this issue, we chose to use the tertiary structures predicted by AlphaFold2 for the protein sequences annotated in UniProt database (39), and create different training, validation, and test sets from them (see Materials and Methods). This approach helps mitigate potential biases arising from variations in experimental techniques and coordinate referencing systems, ensuring a more consistent and standardized dataset for training our model.

Protein design is a multifaceted problem in which many different design task can be considered(16; 17; 7; 18). Multiple strategies have been developed to address these diverse design modalities. However, some of these strategies require a high level of technical knowledge, are time and cost-intensive, or are restricted to a specific problem. On its side, in the field of NLP the GPT series successfully faced the challenge of solving different language problems with a unique model (19; 20; 21). This fact inspired us to train a small Transformer with an encoder-decoder structure, similar to the one described in Ashish et al. work (22), which we name StructureGPT. Our model tackles the general task of predicting the amino acid sequence corresponding to a tertiary structure operating in an autoregressive manner. This way StructureGPT can predict the probability of the next amino acid in the sequence (*AA*_*i*_), conditioned on the preceding amino acids (*AA*_*<i*_), i.e., P(*AA*_*i*_|*AA*_*<i*_).

Model training was conducted independently with two datasets generated from different predictions made by AlphaFold2. In both cases (see Table 1), the training concluded with an accuracy of around 92% (Multiclass accuracy over the train dataset), and no overfitting was observed (around 92% Multiclass Accuracy over the validation dataset). A common way to measure and compare the performance of deep learning models in the field of protein design is the native sequence recovery rate (NSR) (3). Considering how the Multiclass accuracy function is computed, the observed accuracy on the validation dataset at the end of training corresponds to the average NSR achieved by the model.

**Table 1.** Multiclass accuracy over training, validation and test sets. The small multiclass accuracy differences between training and validation datasets were interpreted as no overfitting occurred during training.

| Dataset | Multiclass accuracy (%) |  |  |
| --- | --- | --- | --- |
|  | Training | Validation | Test |
| SwissProtDB | 91.87 | 91.86 | 91.87 |
| AlphaFoldDB_1M | 92.13 | 91.88 | 91.90 |

### 3.3 Testing StructureGPT generation

To illustrate the utility of StructureGPT for addressing multiple design tasks, in this section we present a detailed description of the main generative capabilities we have observed in the model and discuss its utility.

The common starting point for solving any design problem with StructureGPT is to leverage the decoder’s ability to generate amino acid sequences. These sequences are conditioned by the structural information that the encoder has received. One of the simplest strategies that can be adopted during the structure-to-sequence translation task is the so-called ‘greedy decode’ strategy, in which the decoder adds to the generated sequence the element with the highest predicted probability of appearing in that position given the elements that precede it. Figure S2 shows the alignments of the original and generated sequences for a group of 5 proteins from the test set randomly chosen. Consistent with the high accuracy obtained during model training (see Material and Methods), it can be seen how the translation of 4 out of the 5 structures leads to sequences that align with 100% identity with the real sequence. In the case of the protein with UniProt ID A0A024R363, it is observed that real and translated sequences share lower identity (99.51%). A detailed examination of the errors made by the model reveals that it inserted an additional aspartic acid in position 249, which causes a mismatch over almost the rest of the sequence. We have observed that sometimes the model also makes invertions between amino acids. It is noteworthy that sometimes insertions precede inversions and vice versa. Given the autoregressive scheme followed by StructureGPT, the insertion errors may explain the appearance of inversions and the same on the contrary case. Inversions, insertions, and deletions are the most frequent errors we observed.

### 3.4 Thermodynamic stability modulation of human frataxin

An alternative strategy for sequence generation, which we have called ‘mutational decode’, is useful for certain design tasks. This strategy consists of considering more than one amino acid class for each position of the sequence generated during translation. Figure S3 shows the alignment of some of the sequences obtained by applying this strategy to mature human frataxin (amino acids 90 to 210). To generate these sequences, the model was asked to consider the two most probable amino acid classes for a set of 14 mutational hotspots found through a conservation analysis (see Materials and Methods). Consistent with the autoregressive generation scheme, the generated sequences had lengths greater, equal to, or less than the original sequence.

A problem that limits the usefulness of these sequences is knowing in advance the physicochemical properties and the biological function of the structure that corresponds them. However, sometimes there is prior information related to the design task that allows filtering these sequences. An example of this idea could be the search for point mutations that lead to thermodynamic stabilization. In such cases, those sequences that have the same length as the original sequence and include mutations only at desired points could be considered as potential solutions. Following these ideas, those mutant versions of the frataxin sequence that had the same length as the original and only included mutations at the hotspots were filtered, thereby obtaining 56 sequences (Figure S4). Next, the effect on the thermodynamic stability of the mutations introduced in each sequence was evaluated using the empiric force field FoldX (25) (see Materials and Methods). Table 2 shows the values of the changes in free energy of folding, Δ*G*_*F*_ , the differences with respect to wild-type (WT) human frataxin, ΔΔ*G*_*F*_ , and the expected biological effect for some of the mutant sequences obtained. It can be seen how FoldX predicts a stabilizing effect for some of the mutants, while for others the predicted effect is destabilizing. A comparative analysis of the stability values observed for the different mutant forms suggests that the T102A mutation on the mature frataxin sequence could have a strongly stabilizing character, while the L97R mutation could cause strong destabilization. It is worth noting that all Δ*G*_*F*_ values calculated were positive, probably because human frataxin is a mitochondrial protein and the calculation was performed considering a temperature of 298 K, which is a temperature severely different from that found in the mitochondrion (32). Also, despite been a powerful tool for calculating the energy difference between two well defined structures, FoldX sometimes leads to false positives (27; 25). Due to all this, the actual effect of these mutations should be elucidated through a complementary experimental analysis. Furthermore, an analysis of the frequencies with which StructureGPT introduced the different mutations (Figure S4) revealed that the frequency of the T102A mutation was around 42.85% despite expected to be stabilizing, while the frequency of the L97R mutation was around 58.92% despite expected to be destabilizing. This fact is consistent with the idea that the model is not specifically trained to enhance stability.

**Table 2.**
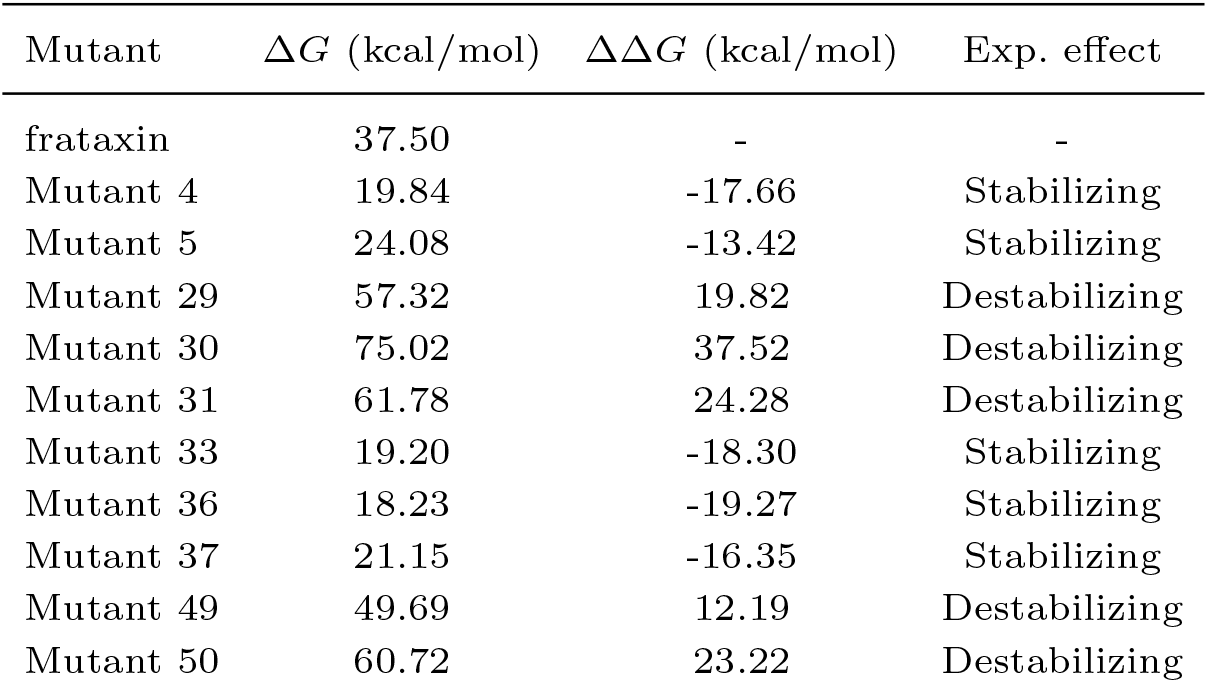
Δ*G*, ΔΔ*G* and the expected biological effect upon mutation calculated for human frataxin and 10 mutants.

### 3.5 Solubility modulation of recombinant human granulocyte-macrophage colony-stimulating factor (rhGM-CSTF)

Another design task that can be addressed using the mutational decode strategy is modifying protein solubility. Similar to modulating stability, in the case of modulating solubility, the first step is to identify those points in the sequence that are susceptible to mutations and contribute poorly, or even negatively, to solubility. Similar to the case of stability modulation, sequences that have the same length as the original sequence and include mutations at desired points are potential solutions to the problem. Figure 3 shows the structure, the intrinsic solubility profiles and structure-corrected intrinsic solubility profiles of rhGM-CSF and 2 of the 72 mutant variants obtained through a similar analysis as carried out with human frataxin (see Materials and Methods). All intrinsic profiles were calculated using the CamSol algorithm (26). It can be observed in the structure-corrected profile of rhGM-CSF (Figure 3B) that isoleucine at position 96 (ILE96) and valine at position 112 (VAL112) contribute negatively to solubility and generate low solubility patches in the protein structure. These patches are formed by glutamine 95 (GLN95) and ILE96, and leuncine 111 (LEU111) and VAL112 respectively. Since all amino acids are exposed on the surface (Figure 3A), they may constitute aggregation hotspots. It can be seen (Figures 3C and 3D) how in the structure-corrected profiles of the two mutant versions of rhGM-CSF, both patches of low solubility have been eliminated. The calculation of the overall solubility, performed as described in (26), on the WT form and the two mutant variants (Table S3) demonstrates how solubility of both mutants has also increased compared to that of the WT form.

**Fig. 3.**
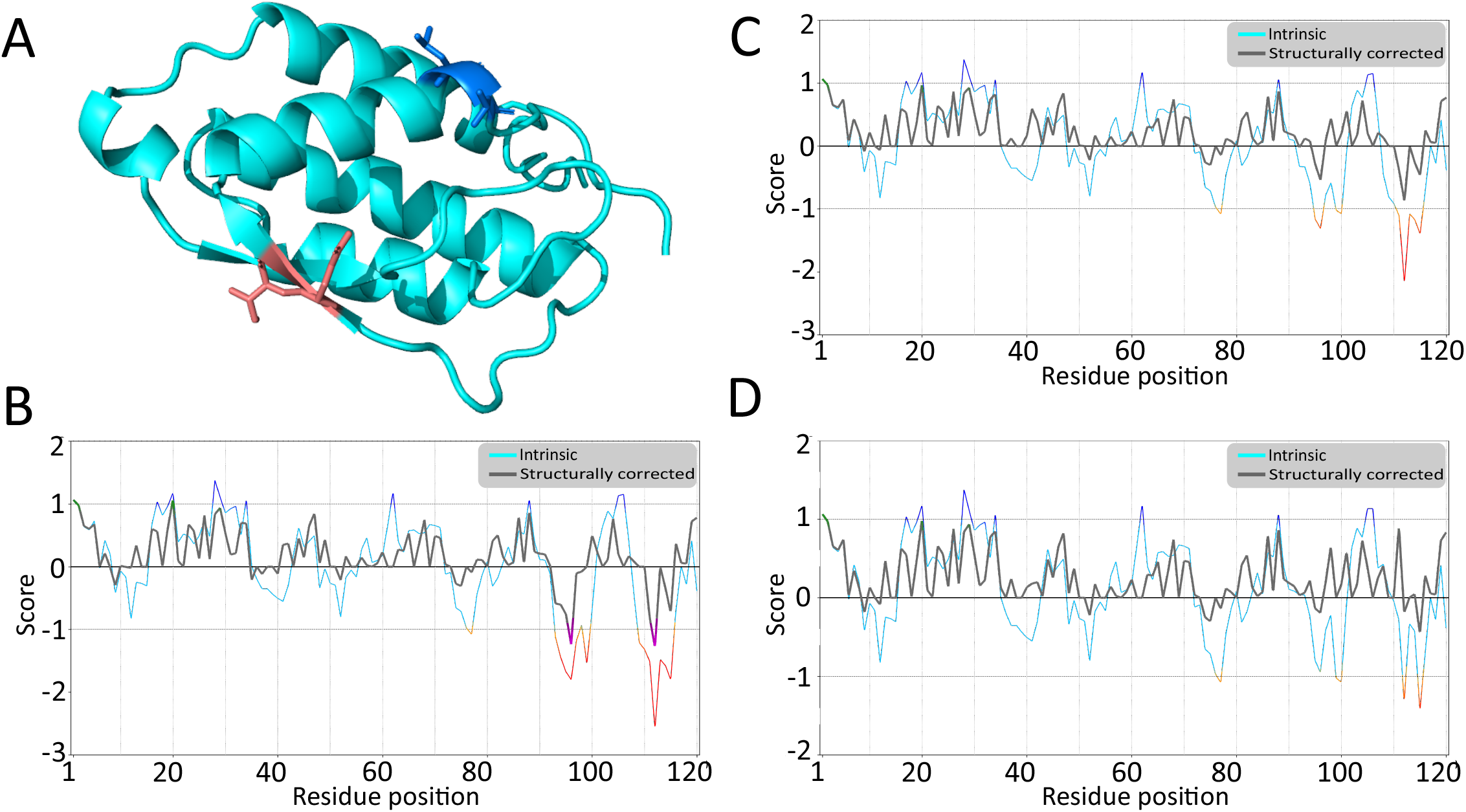
Modification of rhGM-CSF solubility with StructureGPT. (A) Structure of rhGM-CSF. The amino acids GLN95 and ILE96 (red), and LEU111 and VAL112 (dark blue) are highlighted on the structure. Intrinsic solubility profile and structure-corrected intrinsic solubility profile of rhGM-CSF (B), mutant 116 (C), and mutant 131 (D). In (B), (C) and (D), intrinsic solubility profiles correspond to cyan curves, whereas structure-corrected intrinsic profiles correspond to grey curves. In structure-corrected intrinsic profiles, regions with low solubility are highlighted in purple, while regions with high solubility are highlighted in green. It can be seen how in both mutant versions the solubility of the two poorly soluble regions has increased.

### 3.6 Inpainting capabilities over human ATP synthase

Sometimes the simplest design strategy might be first to design a small portion of the structure and subsequently search for the amino acid sequence that stabilizes it. Drawing a parallel with NLP, this would be equivalent to selecting a group of words as a starting point and then seeking a meaningful sentence containing those words. By choosing the tertiary structure as a starting point and representing it sequentially, with the amino acid as the unit of the sequence, we aim to liken the structure-sequence translation to a problem of translating between two languages. Thus, it might be possible to leverage the generative capabilities of LM to incorporate the missing information in the designed structure portion during the translation process.

Similar to NLP models, one of the most remarkable abilities of StructureGPT is its capability to perform ‘inpainting’, that is, replenishing missing information during the task of translating protein structures into sequences. Figure 4 illustrates the overlay of human ATP synthase subunit A structures in its wild-type form alongside two variants derived using an inpainting strategy (see Material and Methods). A darker region representing the sequence segment (amino acids 158 to 198) restored by StructureGPT during translation is highlighted on each structure. Despite variations in the sequences obtained (Figure S7), the structures of each variant maintain a high degree of similarity.

**Fig. 4.**
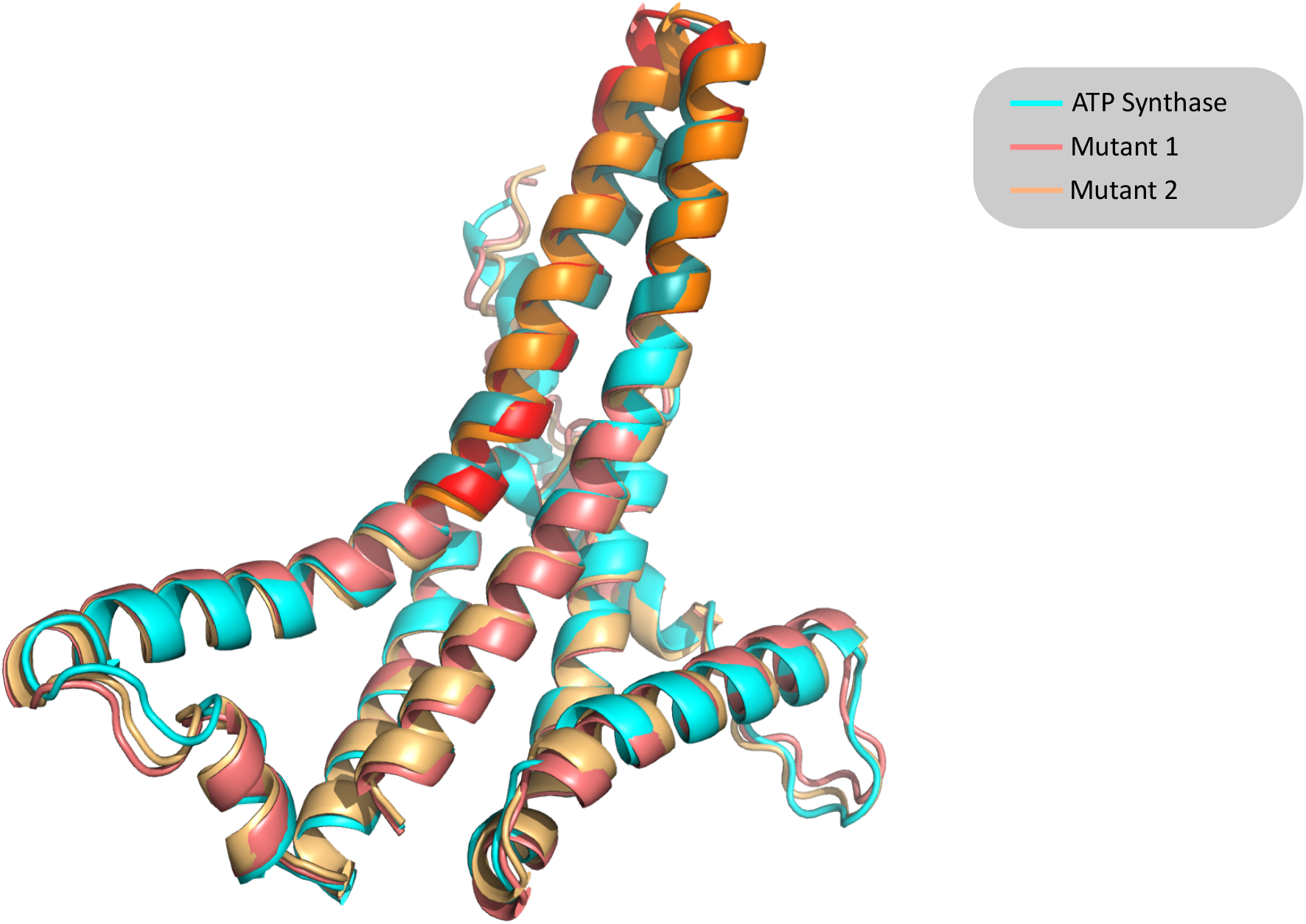
Structure overlay of inpainted ATP synthase variants. Overlay of the 3D structures of the WT human ATP synthase subunit A (cyan) and two mutant variants (red and orange) obtained through the inpainting strategy. The original part of the structure (gray) and the corresponding redesigned parts (dark orange and dark red) are highlighted on all three structures. These parts correspond to the amino acids between positions 158 and 198 of the original protein sequence.

To quantify the differences between these sequences, each was subjected to multiple sequence alignments against the UniRef90 database. Subsequent analyses calculated sequence identity percentages between each reference sequence and the rest of sequences in the alignment. The distributions of identity values obtained were then used to estimate differences between means. Similarly, sequence identity percentages considering only the inpainted region were calculated and analyzed in the same manner. Figure 5 displays the distributions of identity percentages extracted from MSA for the two variants ant the ATP synthase considering both, the complete sequences and only the inpainted regions. Results show significant differences in mean identities across alignments for both full sequences and inpainted regions (p-value *<*0.001). Given that the sequences of the three ATP synthase variants differ only in the inpainted region (Figure S7), this highlights two key points: a significant alteration of alignments for each sequence and that one source of observed variability in alignments stems from changes introduced in the inpainted region.

**Fig. 5.**
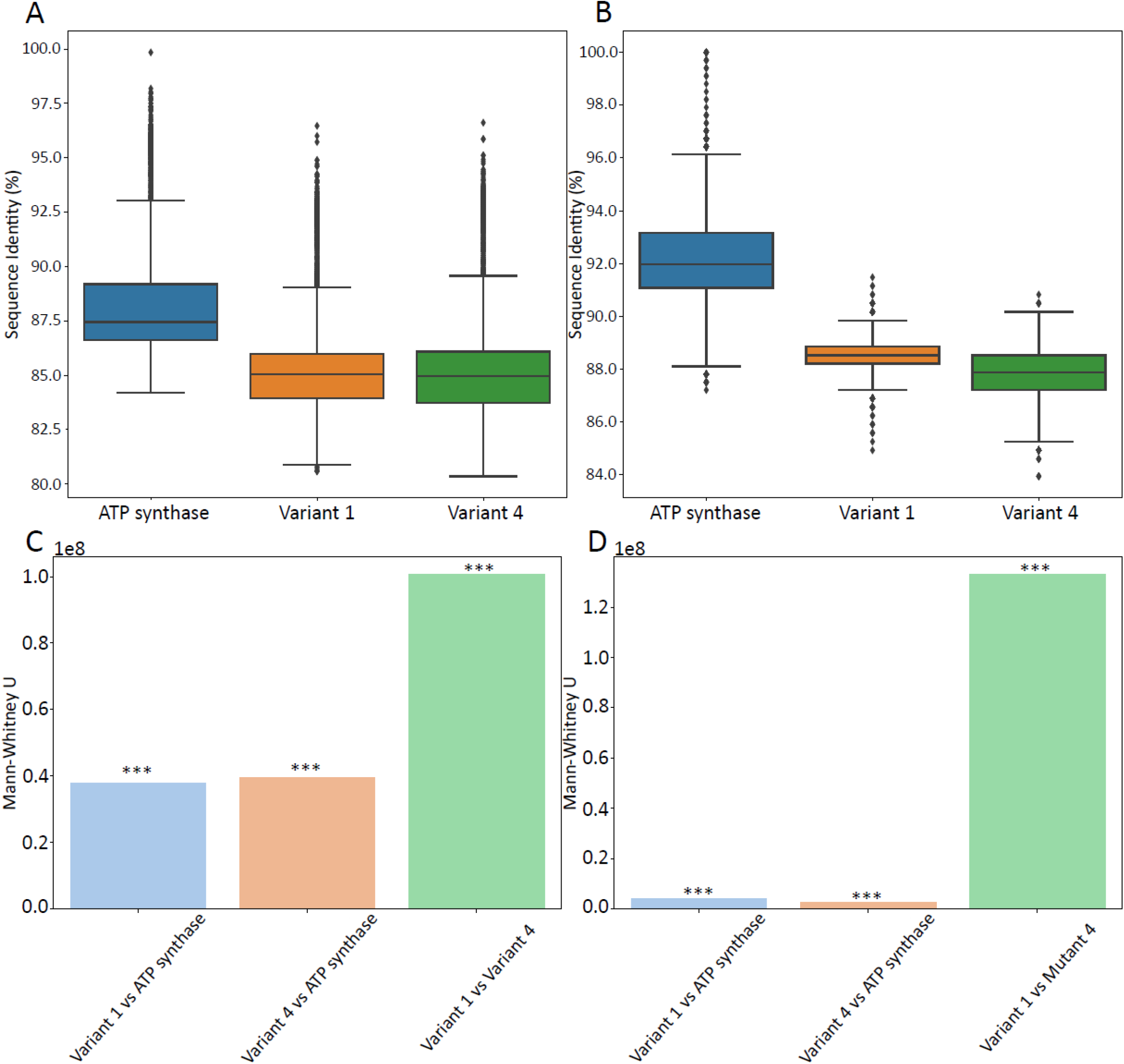
Quantitative analysis of inpainted ATP synthase variants. (A) Distributions of sequence identity percentages from the multiple sequence alignment (MSA) for the complete sequences of human ATP synthase subunit A (blue), variant 1 (orange), and variant 4 (green). (B) Distributions of sequence identity percentages from the MSA of inpainted regions for human ATP synthase (blue), variant 1 (orange), and variant 4 (green). (C) Mann-Whitney U statistical analysis comparing mean sequence identities between ATP synthase and variant 1 (blue), ATP synthase and variant 4 (orange), and variant 1 versus variant 4 (green) for whole sequences. (D) Mann-Whitney U statistical analysis comparing mean sequence identities for the inpainted regions between ATP synthase and variant 1 (blue), ATP synthase and variant 4 (orange), and variant 1 versus variant 4 (green). In (C) and (D) statistical significance is denoted by asterisks (***), with a p-value *<*0.001, highlighting the inpainted region as a primary source of variability in the alignments.

Concurrently, the structures for each variant were calculated using AlphaFold2. Table 3 shows the root mean square deviation, (RMSD_*mc*_), global distance test total score, (GDT-TS_*mc*_), global distance test high accuracy, (GDT-HA4_*mc*_) and local distance difference test for main chain atoms, (lDDT_*mc*_), obtained from structural alignments. Collectively, these metrics suggest that all three structures are highly similar to each other, with the redesigned variants showing greater similarity to each other than to the original ATP synthase. This finding supports the idea that StructureGPT can accurately comprehend three-dimensional protein structures and, consequently, generate sequences that lead to similar structures, even if they differ from the original protein sequence. Likely, the modifications introduced by StructureGPT induce functional changes in ATP synthase, although further experimental analysis is required to ascertain the nature of these changes.

**Table 3.**
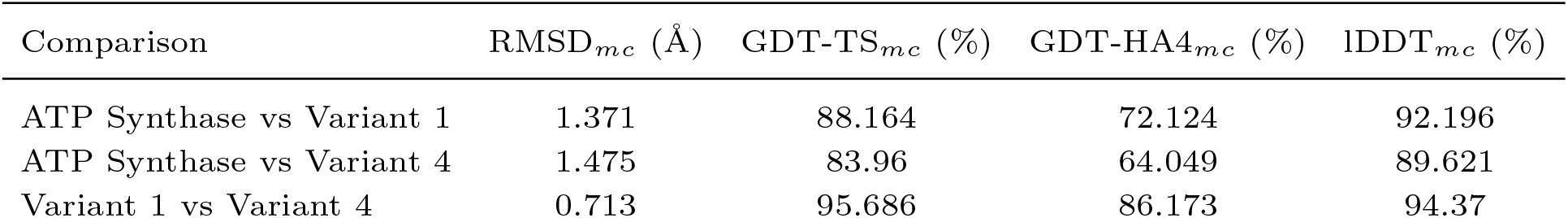
Structural comparison between human ATP synthase and inpainted variants. RMSD_*mc*_, GDT-TS_*mc*_, and GDT-HA4_*mc*_ calculated for the alignment of ATP synthase, Variant 1 and Variant 4 structures are shown.

## 4. Conclusion

We introduced StructureGPT, an innovative deep learning model designed to translate protein tertiary structures into corresponding amino acid sequences using a novel encoding system. By leveraging the parallels between protein structures and human languages, StructureGPT autoregressively generates amino acid sequences, enhancing our ability to design proteins with specific functionalities. Key to our methodology was the development of a unique atom encoding system that effectively captures the spatial and contextual information of amino acids in protein structures. This encoding not only facilitated the training of our model but also ensured that the generated sequences retained functional relevance to their structural inputs. The training datasets used provided a standardized and robust foundation for developing our model, ensuring consistency across training and validation phases. StructureGPT’s generative capabilities were particularly demonstrated in its application to enhance solubility and stability—two critical attributes that often limit the practical use of proteins in industrial and pharmaceutical contexts. By exploiting the model’s generative capabilities, we were able to propose modifications to protein sequences that are predicted to improve these properties. Additionally, the inpainting capabilities of StructureGPT showcased its understanding of protein tertiary structure, enabling the generation of diverse protein sequences that, despite their differences, led to structurally similar outcomes. However, StructureGPT presents certain limitations that delineate areas for future research and development. Currently, the model does not handle quaternary protein structures, which are crucial for understanding complex biological functions that involve multiple subunits interacting. Additionally, direct prediction of functional outcomes from structure-derived sequences remains an unexplored territory that holds significant potential for expanding the utility of StructureGPT in functional genomics and proteomics.

## Author contributions statement

N.Z. designed research, N.Z. and P.U. performed research, N.Z. and P.U. analyzed data, N.Z. and H.B. wrote the paper

## Acknowledgments

We thank Dr. Javier Fumanal and Iosu Rodríguez for their comments and advices. We thank ChatGPT for helping in the translation of this manuscript. We thank Navarra de Servicios y Tecnologías S.A. (NASERTIC) for providing the computation resources. We thank Navarra Artificial Intelligence Research (NAIR) Center. This work was supported by grants and funds from Fundación ONCE, the European Social Fund (ESF) and the Spanish Ministry of Science (project PID2022-136627NB-I00).

---

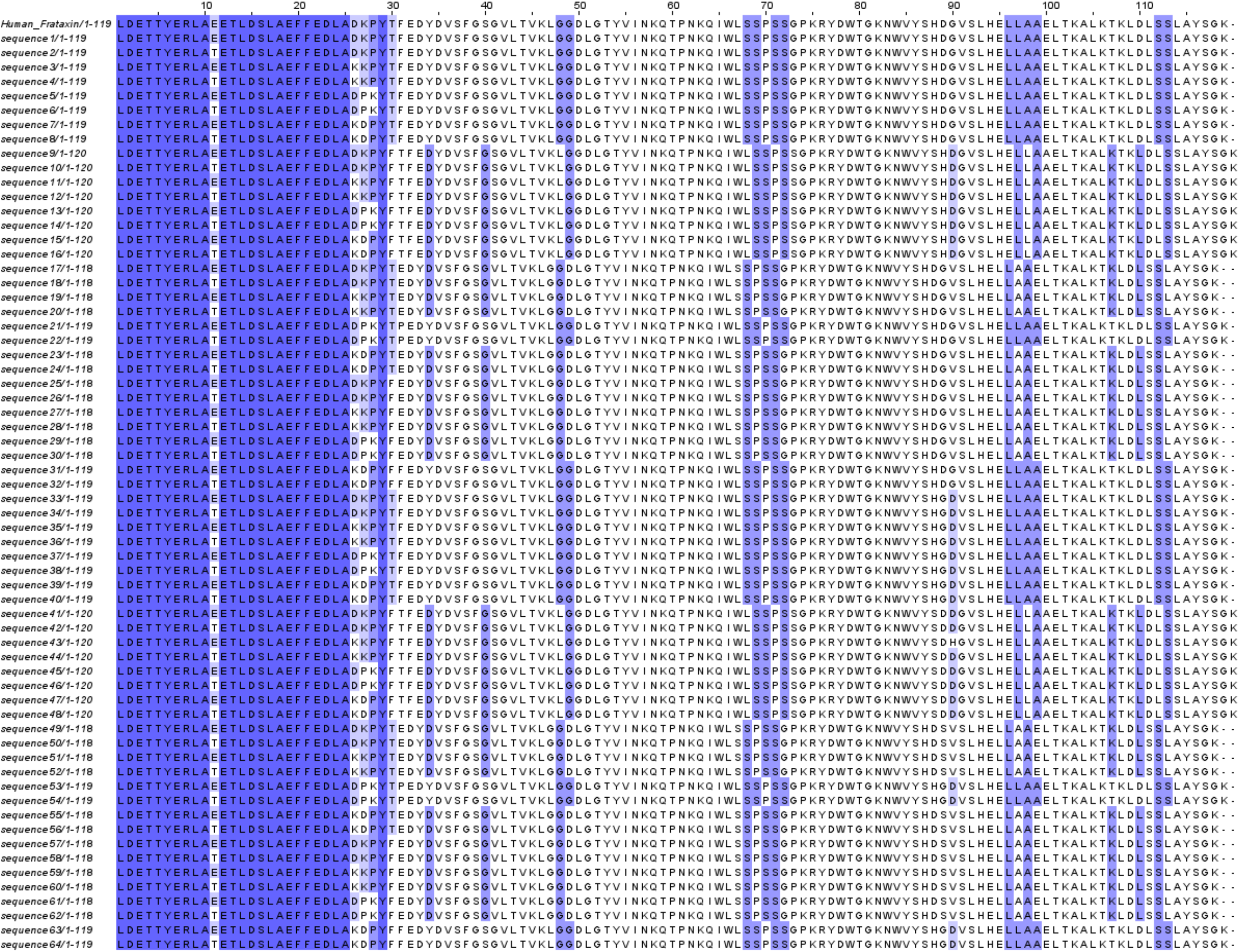

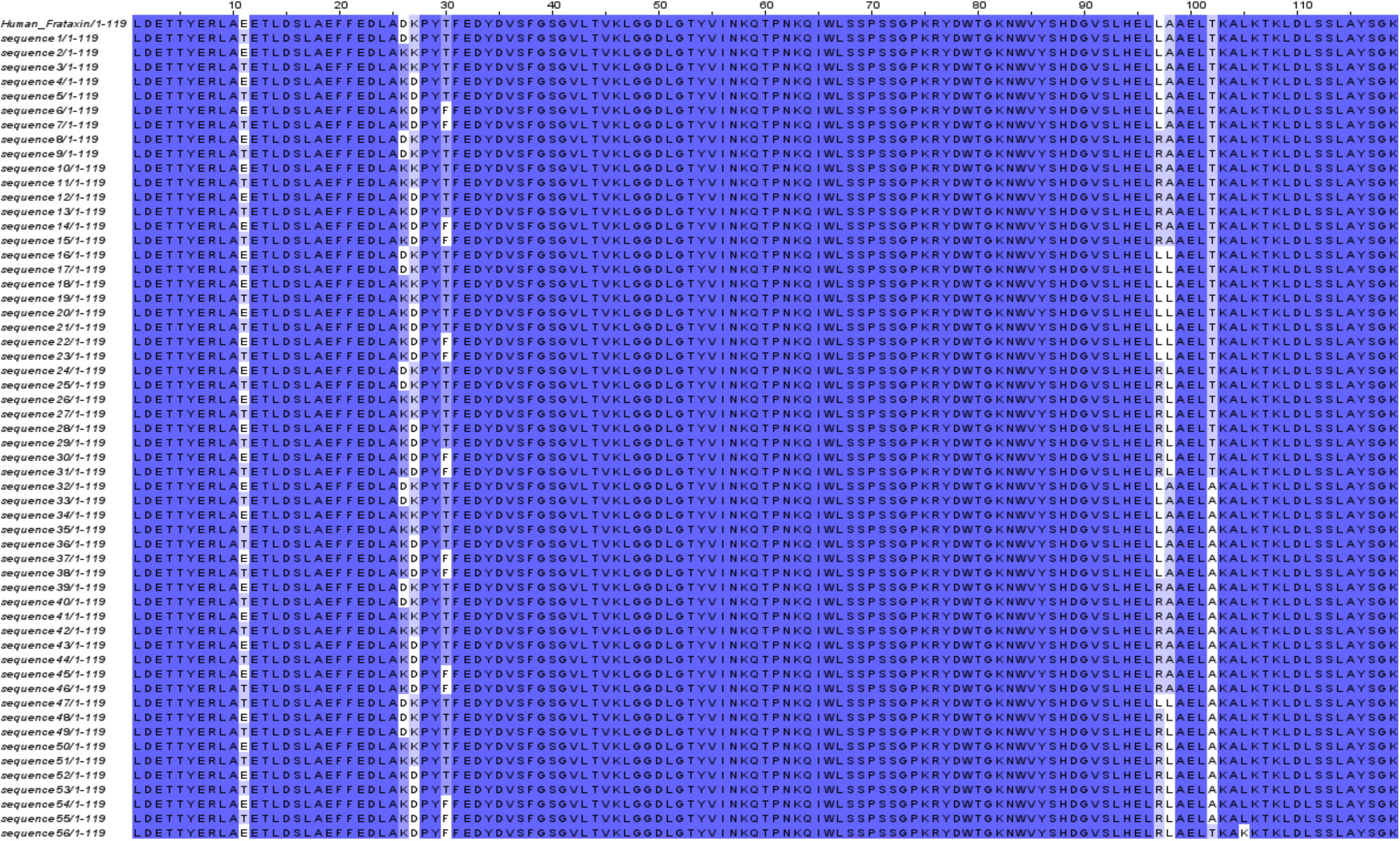

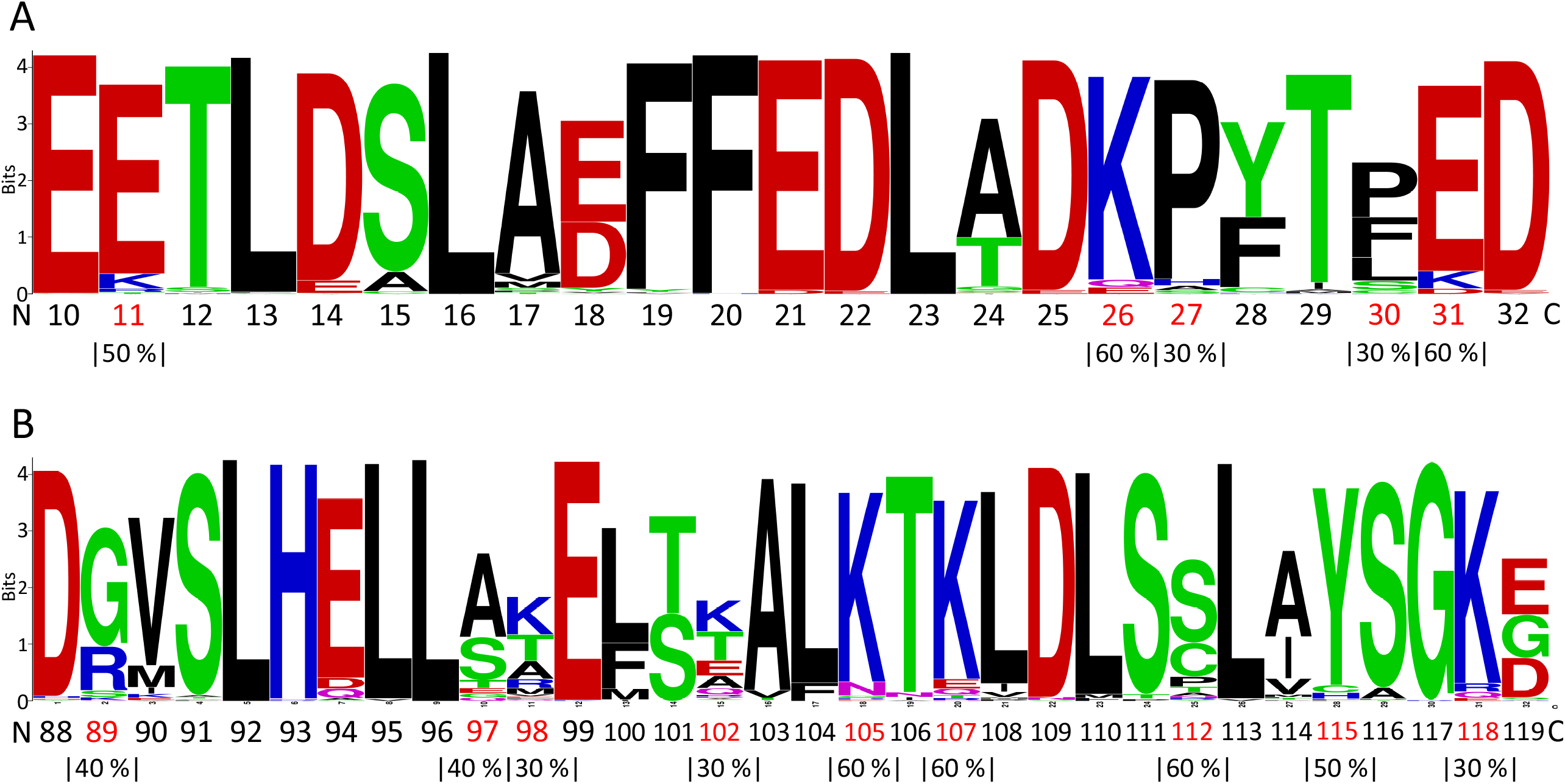

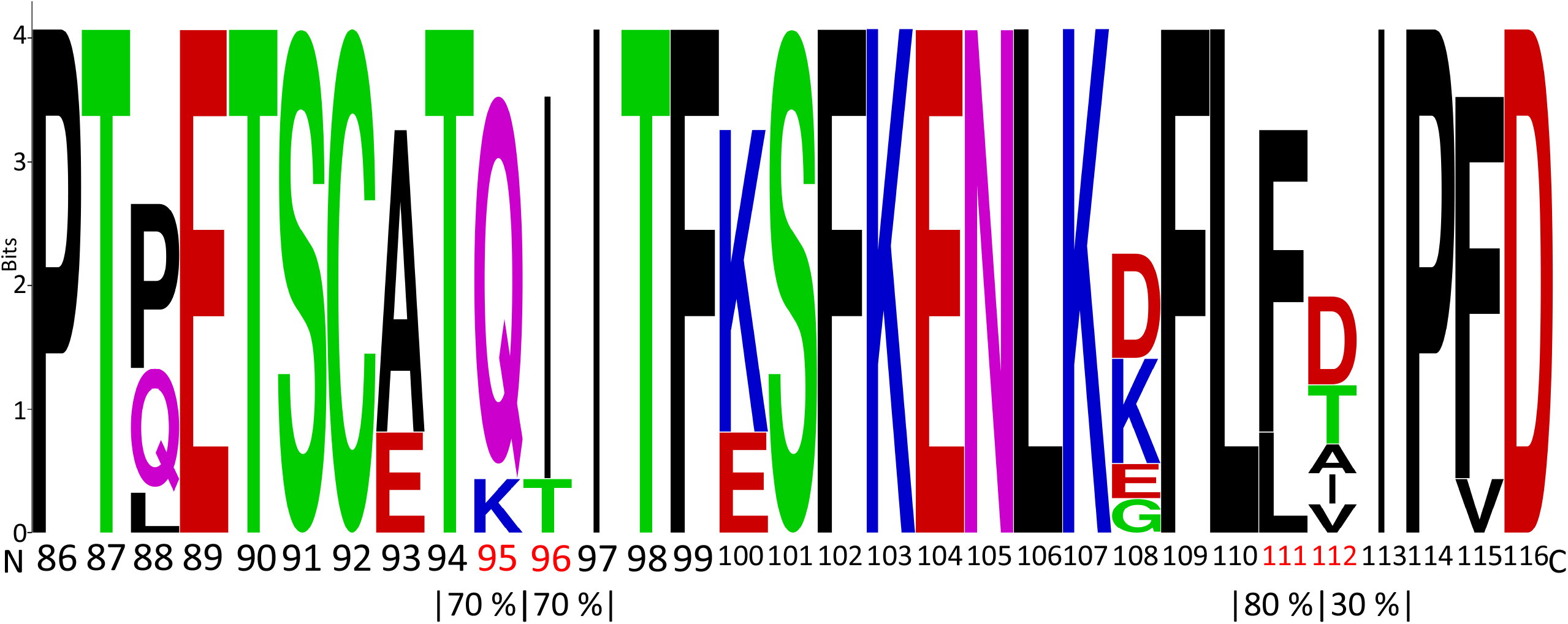

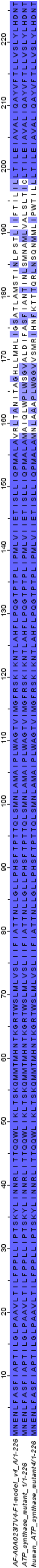

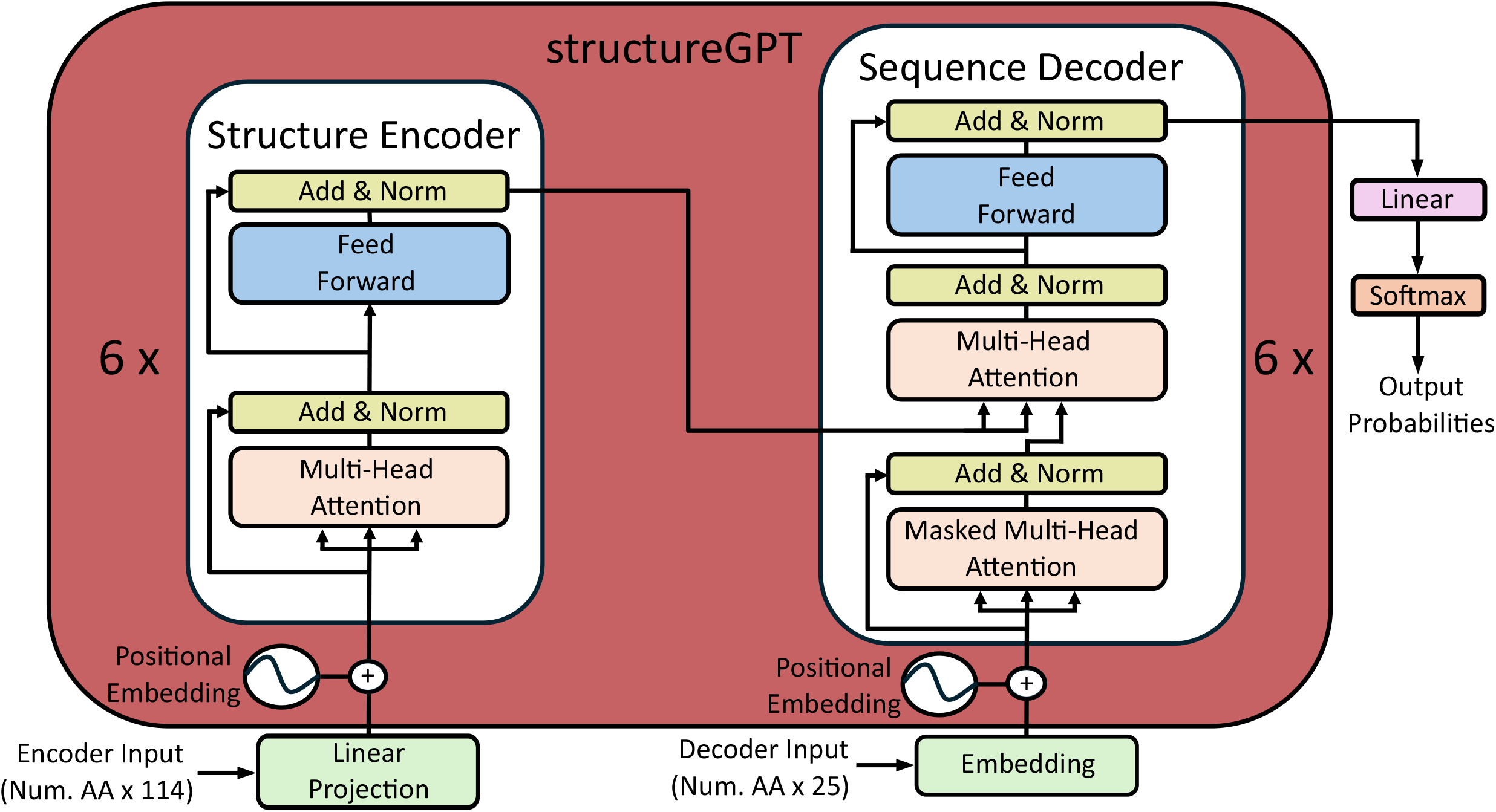

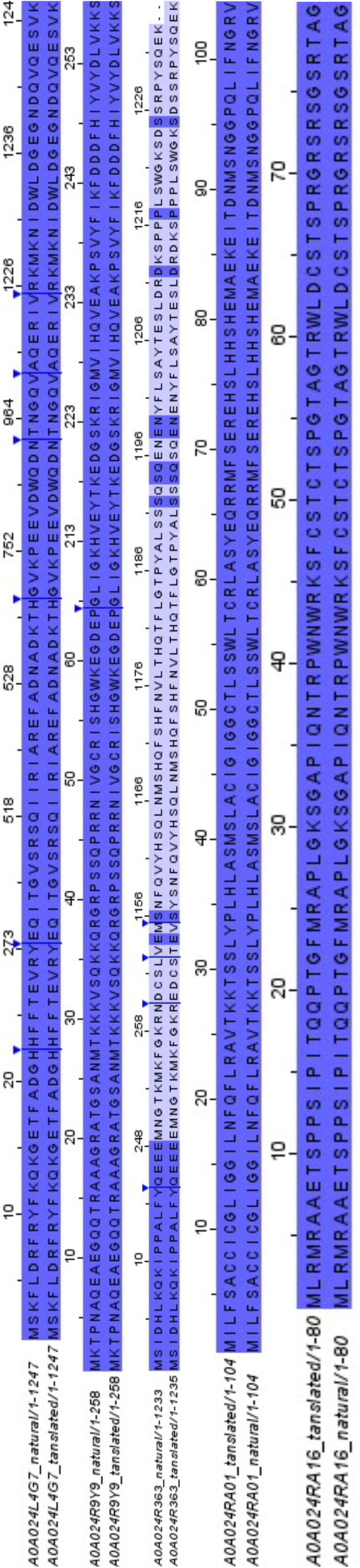

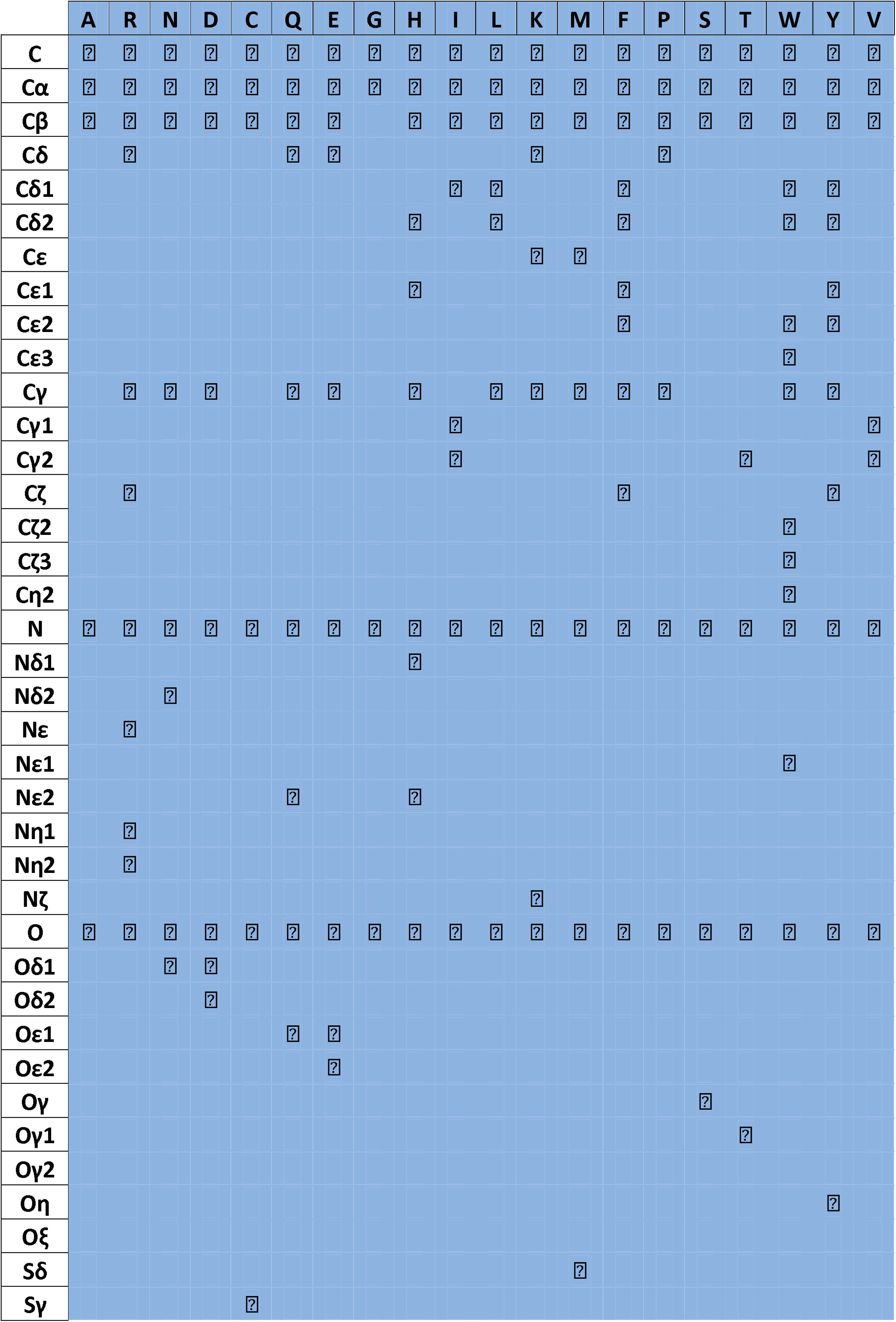

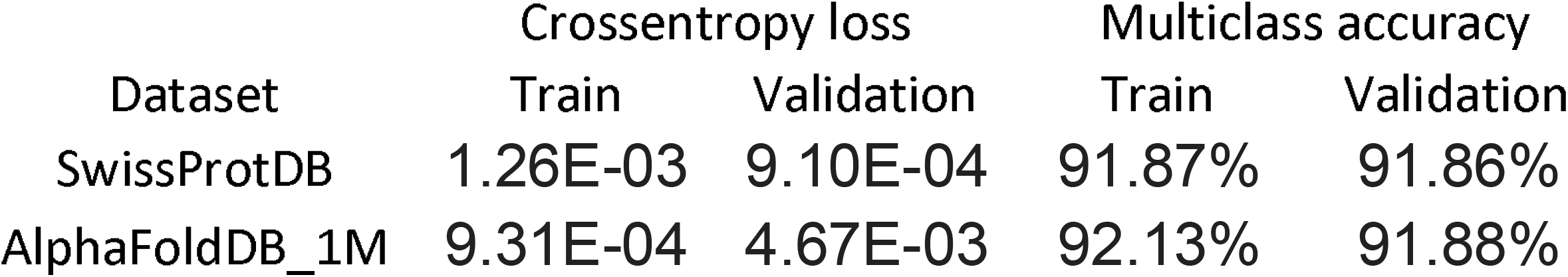

| Dataset | Crossentropy loss |  | Multiclass accuracy |  |
| --- | --- | --- | --- | --- |
|  | Train | Validation | Train | Validation |
| SwissProtDB | 1.26E-03 | 9.10E-04 | 91.87% | 91.86% |
| AlphaFoldDB_1M | 9.31E-04 | 4.67E-03 | 92.13% | 91.88% |

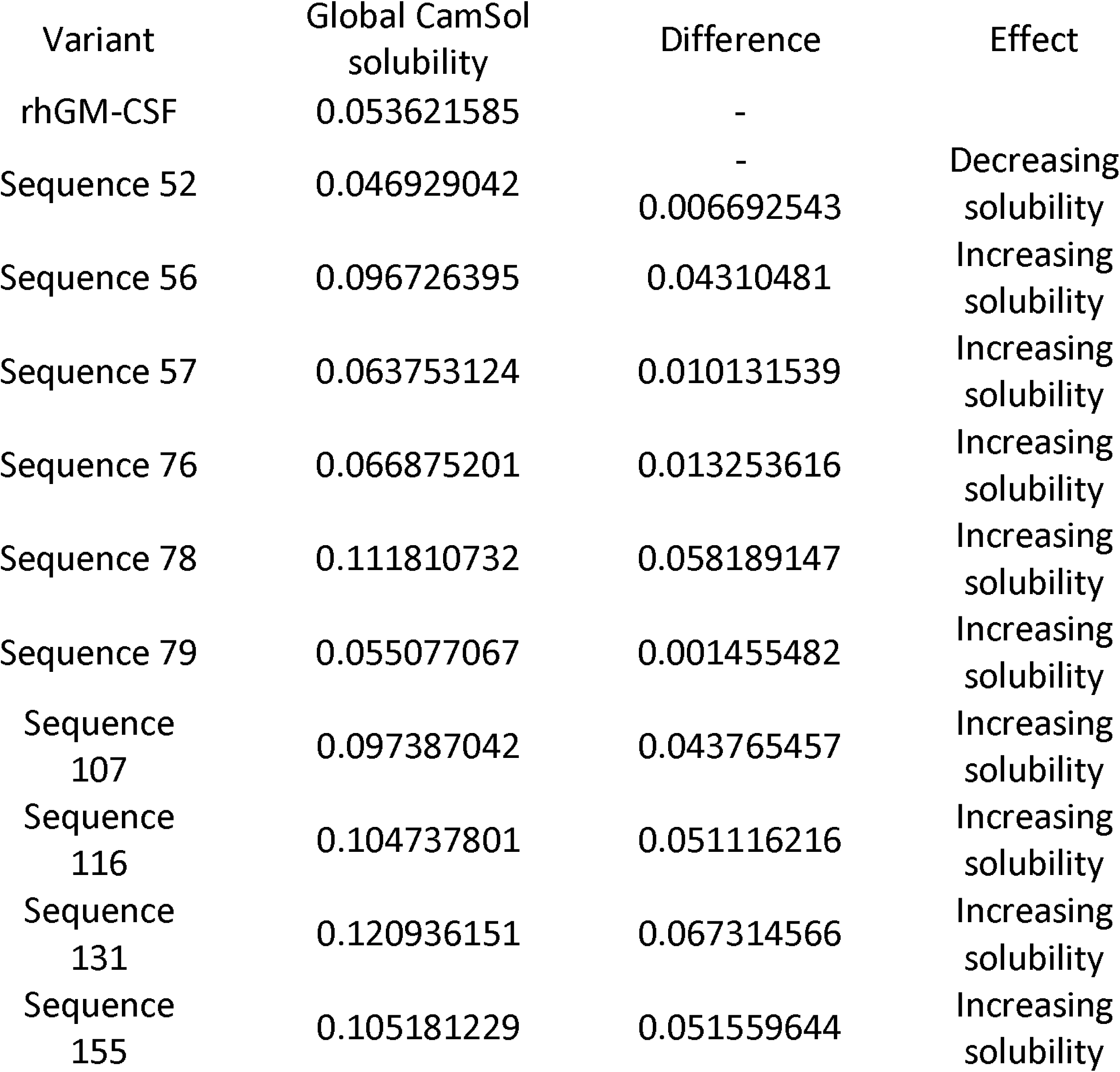

