## Supporting Information for "Protein Design with StructureGPT: a Deep Learning Model for Protein Structure-to-Sequence Translation"

**Table S1. Atomic classes for the atom encoding.** The presence in each of the 20 naturally occurring amino acids of 37 out of the 38 atomic classes found in the structure of proteins is shown. Atomic class 38 mentioned in the main text corresponds to the terminal oxygen.

| | A | R | N | D | C | Q | E | G | H | I | L | K | M | F | P | S | T | W | Y | V |
|---|---|---|---|---|---|---|---|---|---|---|---|---|---|---|---|---|---|---|---|---|
| **C** | ✓ | ✓ | ✓ | ✓ | ✓ | ✓ | ✓ | ✓ | ✓ | ✓ | ✓ | ✓ | ✓ | ✓ | ✓ | ✓ | ✓ | ✓ | ✓ | ✓ |
| **Cα** | ✓ | ✓ | ✓ | ✓ | ✓ | ✓ | ✓ | ✓ | ✓ | ✓ | ✓ | ✓ | ✓ | ✓ | ✓ | ✓ | ✓ | ✓ | ✓ | ✓ |
| **Cβ** | ✓ | ✓ | ✓ | ✓ | ✓ | ✓ | ✓ | | ✓ | ✓ | ✓ | ✓ | ✓ | ✓ | ✓ | ✓ | ✓ | ✓ | ✓ | ✓ |
| **Cδ** | | ✓ | | | | ✓ | ✓ | | | | | ✓ | | | ✓ | | | | | |
| **Cδ1** | | | | | | | | | | ✓ | ✓ | | | ✓ | | | | ✓ | ✓ | |
| **Cδ2** | | | | | | | | | ✓ | | ✓ | | | ✓ | | | | ✓ | ✓ | |
| **Cε** | | | | | | | | | | | | ✓ | ✓ | | | | | | | |
| **Cε1** | | | | | | | | | ✓ | | | | | ✓ | | | | | ✓ | |
| **Cε2** | | | | | | | | | | | | | | ✓ | | | | ✓ | ✓ | |
| **Cε3** | | | | | | | | | | | | | | | | | | ✓ | | |
| **Cγ** | | ✓ | ✓ | ✓ | | ✓ | ✓ | | ✓ | | ✓ | ✓ | ✓ | ✓ | ✓ | | | ✓ | ✓ | |
| **Cγ1** | | | | | | | | | | ✓ | | | | | | | | | | ✓ |
| **Cγ2** | | | | | | | | | | ✓ | | | | | | | ✓ | | | ✓ |
| **Cζ** | | ✓ | | | | | | | | | | | | ✓ | | | | | ✓ | |
| **Cζ2** | | | | | | | | | | | | | | | | | | ✓ | | |
| **Cζ3** | | | | | | | | | | | | | | | | | | ✓ | | |
| **Cη2** | | | | | | | | | | | | | | | | | | ✓ | | |
| **N** | ✓ | ✓ | ✓ | ✓ | ✓ | ✓ | ✓ | ✓ | ✓ | ✓ | ✓ | ✓ | ✓ | ✓ | ✓ | ✓ | ✓ | ✓ | ✓ | ✓ |
| **Nδ1** | | | | | | | | | ✓ | | | | | | | | | | | |
| **Nδ2** | | | ✓ | | | | | | | | | | | | | | | | | |
| **Nε** | | ✓ | | | | | | | | | | | | | | | | | | |
| **Nε1** | | | | | | | | | | | | | | | | | | ✓ | | |
| **Nε2** | | | | | | ✓ | | | ✓ | | | | | | | | | | | |
| **Nη1** | | ✓ | | | | | | | | | | | | | | | | | | |
| **Nη2** | | ✓ | | | | | | | | | | | | | | | | | | |
| **Nζ** | | | | | | | | | | | | ✓ | | | | | | | | |
| **O** | ✓ | ✓ | ✓ | ✓ | ✓ | ✓ | ✓ | ✓ | ✓ | ✓ | ✓ | ✓ | ✓ | ✓ | ✓ | ✓ | ✓ | ✓ | ✓ | ✓ |
| **Oδ1** | | | ✓ | ✓ | | | | | | | | | | | | | | | | |
| **Oδ2** | | | | ✓ | | | | | | | | | | | | | | | | |
| **Oε1** | | | | | | ✓ | ✓ | | | | | | | | | | | | | |
| **Oε2** | | | | | | | ✓ | | | | | | | | | | | | | |
| **Oγ** | | | | | | | | | | | | | | | | ✓ | | | | |
| **Oγ1** | | | | | | | | | | | | | | | | | ✓ | | | |
| **Oγ2** | | | | | | | | | | | | | | | | | | | | |
| **Oη** | | | | | | | | | | | | | | | | | | | ✓ | |
| **Oξ** | | | | | | | | | | | | | | | | | | | | |
| **Sδ** | | | | | | | | | | | | | ✓ | | | | | | | |
| **Sγ** | | | | | ✓ | | | | | | | | | | | | | | | |

**Table S2. Training and validation accuracies and loss values of StructureGPT.** The Cross-entropy loss and multiclass accuracies developed by StructureGPT over the two datasets considered in this work are shown. In both cases the small accuracy differences observed over training and validation datasets was interpreted as there was no overfitting of the model.

| | Cross-entropy loss | | |
|---|---|---|---|
| **Dataset** | Training | Validation | Test |
| **SwissProtDB** | 1.26E-03 | 9.10E-04 | 5.95E-04 |
| **AlphaFoldDB_1M** | 9.31E-04 | 4.67E-03 | 3.63E-03 |

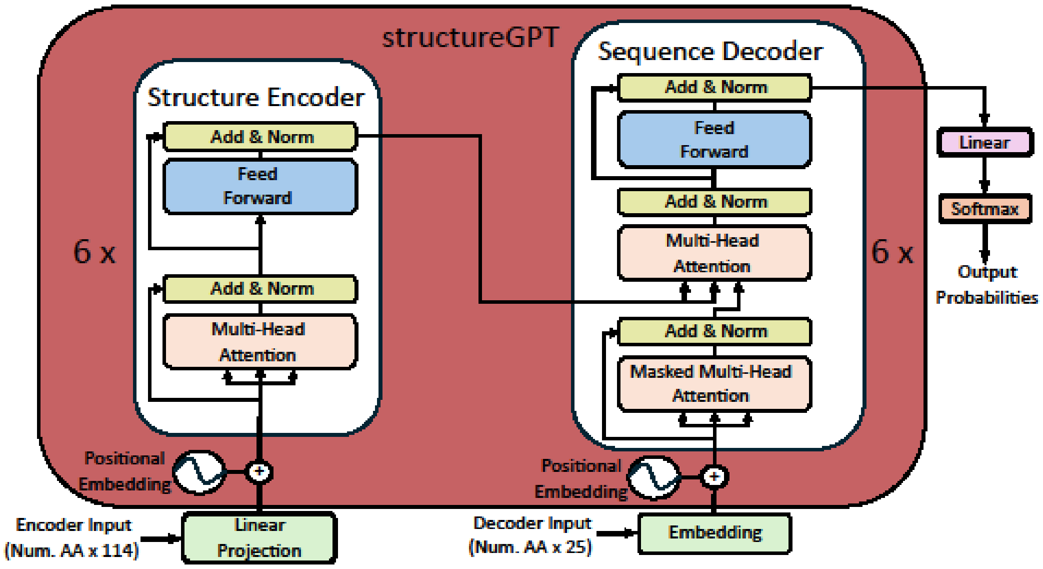


**Fig. S1. Model architecture of StructureGPT.** The overall architecture is composed by two main parts: the encoder and the decoder. The encoder receives and encodes the structure information coming from the atom encoding representation. On the other hand, the decoder receives the encoded structural information, then decodes and converts it into the predicted sequence. Both, the encoder and the decoder of StructureGPT are composed of 6 building blocks, made up by stacks of multi-head attention, batch normalization and linear (feed forward) layers. Previous to entering the enconder, the atom encoded information is projected and enriched with positional encoding information. In a similar way, the information entering the decoder from the outside is embedded and enriched with positional encoding information.

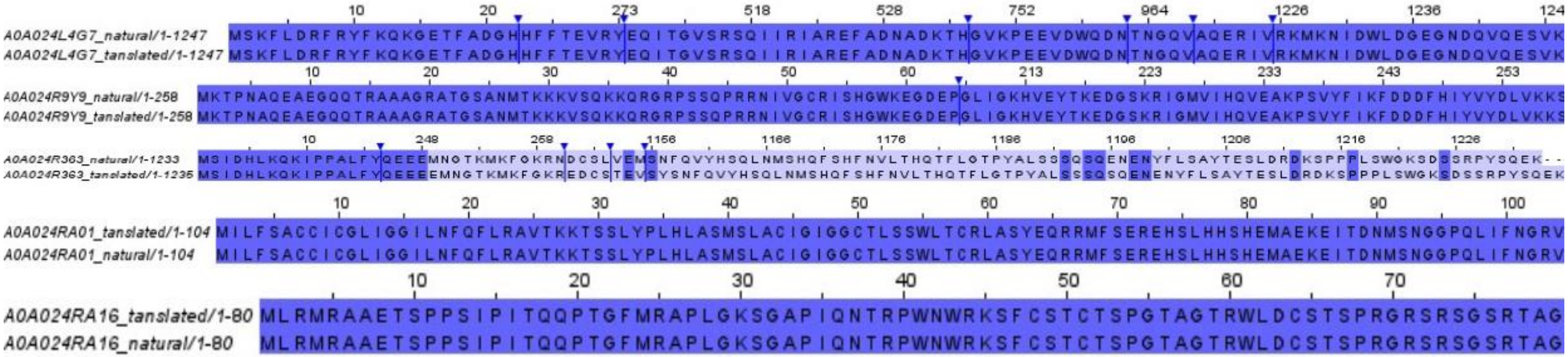


**Fig. S2. Alignment of translated sequences.** From top to bottom, the alignment between natural and translated sequences for nitrate reductase (UniProt ID A0A024L4G7), protein with UniProt ID A0A024R9Y9, protein with UniProt ID A0A024R363, protein with UniProt ID A0A024RA01, and protein with UniProt ID A0A024RA16 are shown. Blue arrows over some of the sequences indicate that some columns are hidden. Aligned sequences are coloured by identity. Darker blue indicates more conserved regions, light blue indicates less conserved regions, and white indicates no conserved regions. The alignment of protein with UniProt ID A0A024R363 shows an insertion in position 249 which causes a subsequent mismatch over almost the whole sequence.

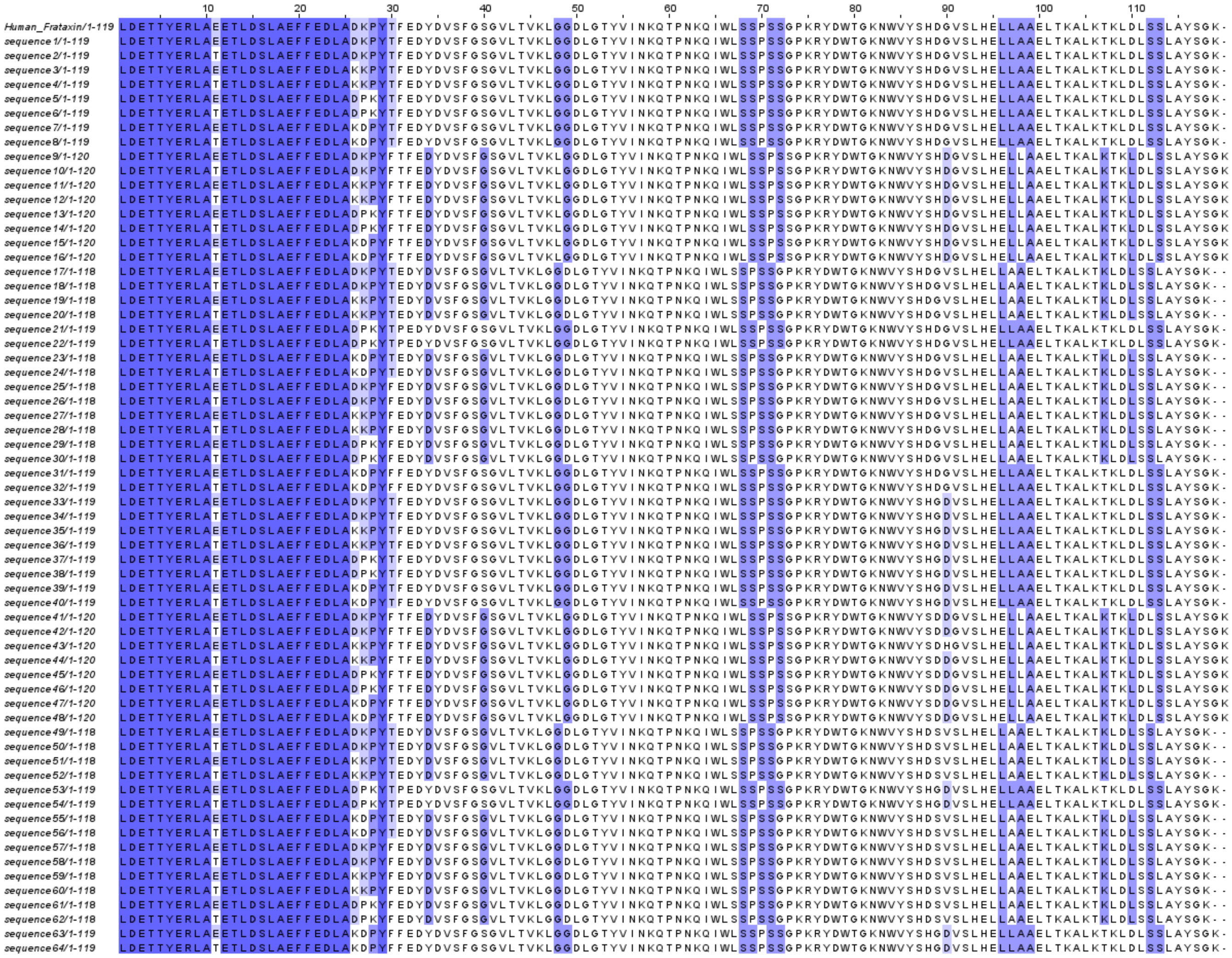

**Fig. S3. Sequence variants of human frataxin.** 64 sequence variants obtained by applying the mutational decode strategy to the mature form of human frataxin (amino acids 90-210, PDB ID 1EKG). The two most probable amino acid classes for each mutational hotspot (see Material and Methods) were considered during translation. Sequences are coloured by identity. Dark blue indicates conserved regions, light blue indicates poorly conserved regions, and white indicates no conservation.

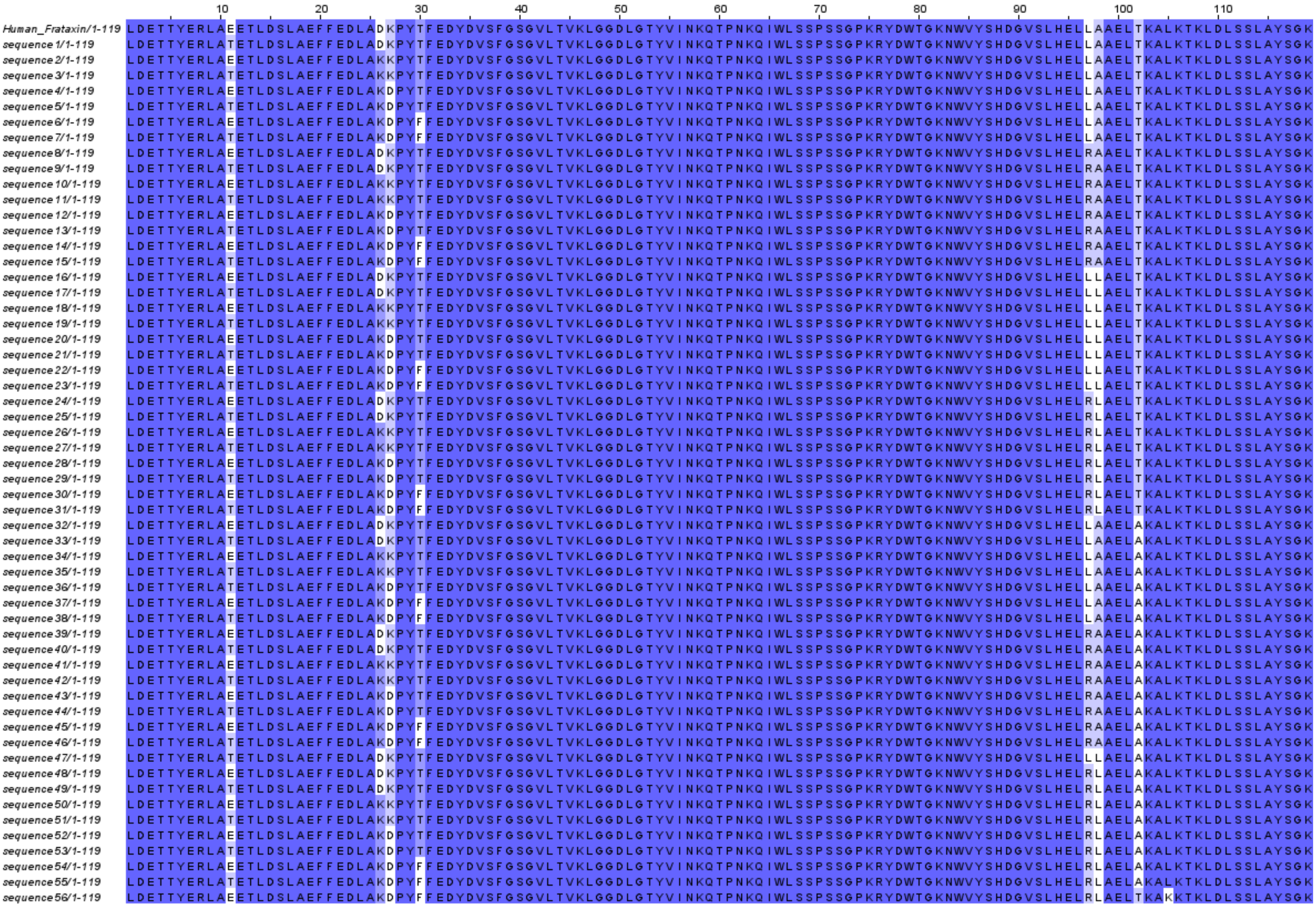

**Fig. S4. Final mature human frataxin mutants.** Mutant variants that match length of the natural human frataxin sequence and include mutations at hotspots identified in the conservation analysis are shown. Sequences are coloured by identity. Dark blue indicates high conservation, light blue indicates poorer conservation, and white indicates no conservation. It can be observed how mutations were included only at some of the hotspots. The frequency of L97R mutation was 58.92% despite expected to be destabilizing, whereas the frequency of T102A mutation was 42.85% despite expected to be stabilizing.

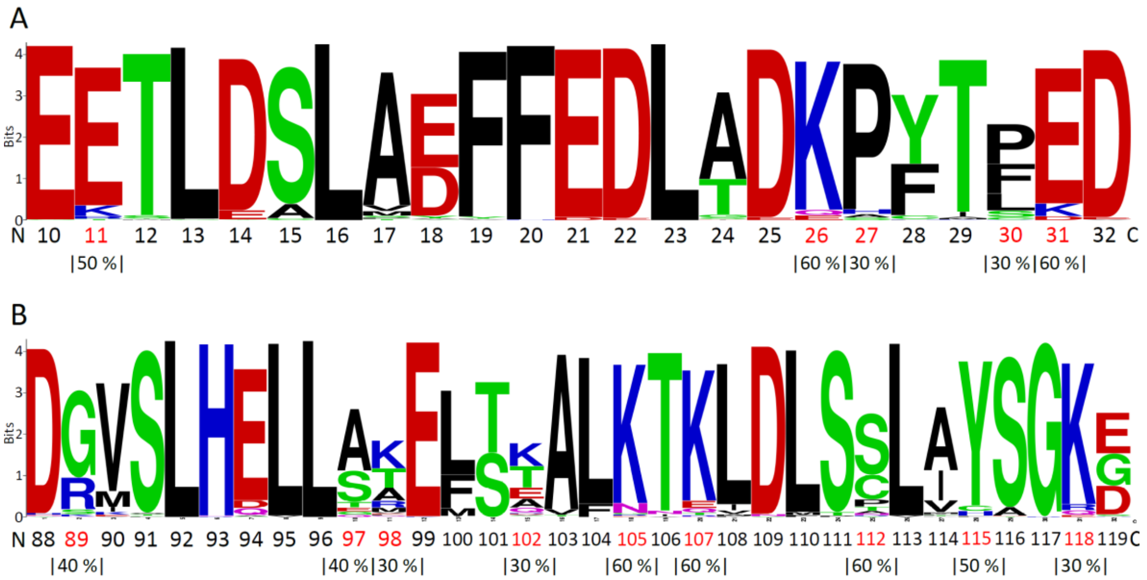


**Fig. S5. Conservation analysis of mature human frataxin.** (A) Conservation analysis for amino acids 10 to 32 of the sequence is shown. (B) Conservation analysis for amino acids 88 to 119 is shown. In both analysis, mutational hotspots are highlighted in red. Beneath each mutational hotspot its conservation percentage is indicated. Acid amino acids are coloured red, basic amino acids are coloured blue, polar amino acids are coloured green, and nonpolar amino acids are coloured black.

**Table S3. Overall solubility of rhGM-CSF and derived mutants.** Overall solubility and solubility differences calculated as described in the main text for 10 out of the 72 mutants obtained are shown. The expected effect is also presented.

| Variant | Global CamSol solubility | Difference | Effect |
|---|---|---|---|
| **rhGM-CSF** | 0.053621585 | - | |
| **Sequence 52** | 0.046929042 | -0.006692543 | Decreasing solubility |
| **Sequence 56** | 0.096726395 | 0.04310481 | Increasing solubility |
| **Sequence 57** | 0.063753124 | 0.010131539 | Increasing solubility |
| **Sequence 76** | 0.066875201 | 0.013253616 | Increasing solubility |
| **Sequence 78** | 0.111810732 | 0.058189147 | Increasing solubility |
| **Sequence 79** | 0.055077067 | 0.001455482 | Increasing solubility |
| **Sequence 107** | 0.097387042 | 0.043765457 | Increasing solubility |
| **Sequence 116** | 0.104737801 | 0.051116216 | Increasing solubility |
| **Sequence 131** | 0.120936151 | 0.067314566 | Increasing solubility |
| **Sequence 155** | 0.105181229 | 0.051559644 | Increasing solubility |

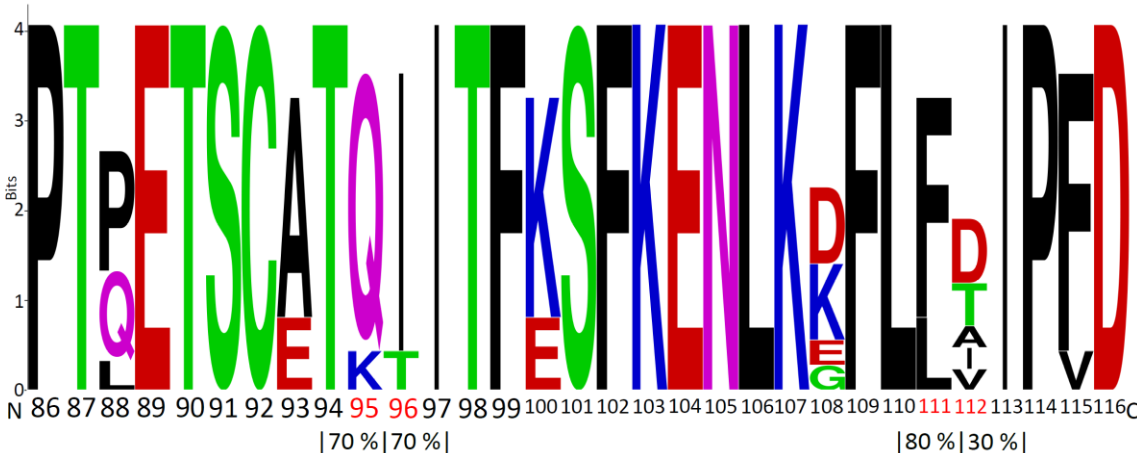


**Fig. S6. Conservation analysis of rhGM-CSF.** Conservation analysis for amino acids 86 to 116. Beneath each hotspot considered for solubility enhancement its conservation percentage is indicated. Acid amino acids are coloured red, basic amino acids are coloured blue, polar amino acids are coloured green, and nonpolar amino acids are coloured black.

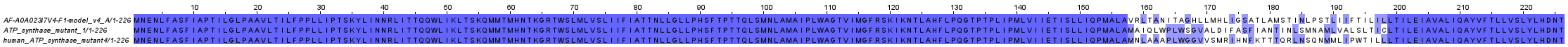


**Fig. S7. Alignment of human ATP synthase and inpainted mutant sequences.** Sequences are coloured by sequence identity with the following code: dark blue for conserver regions, light blue for poorly conserved regions, and white for non-conserved regions. The modified region corresponds to amino acids between valine 158 and leucine 198. It can be seen how only the modified region varies among the two mutants and human ATP synthase.
